# Neuromechanical Justification of Lower-Limb Functional Tests for a Return to Running: A Muscle Coordination Analysis

**DOI:** 10.1101/2023.06.15.545091

**Authors:** Hiroki Saito, Ayu Yamano, Nanae Suzuki, Kazuya Matsushita, Hikaru Yokoyama, Atsushi Sasaki, Tatsuya Takahashi, Kimitaka Nakazawa

## Abstract

**Introduction:** This study aimed to explore the shared muscle synergies between running and functional tests that are commonly used when considering the return to running (RTR) after sports-related injuries. We hypothesized that shared muscle synergies would differ among tasks, providing insights into prioritizing functional tests in the context of RTR.

**Methods:** Ten healthy male participants were recruited to perform running and 9 functional tasks and their 16 lower limb and trunk muscle activities were recorded using electromyography (EMG). Non-negative matrix factorization (NMF) was applied to the collected EMG data to explore shared muscle synergies between running and the functional tasks. We compared the percentages of shared synergies and temporal patterns between running and each functional test.

**Results:** Although all functional tests exhibited shared muscle synergies with running, the walk (75% [40%-100%]), single leg hops with 30% of maximum distance (SLH30) (60% [20%-100%]), and stepup (63% [0%-100%]) tasks displayed significantly higher percentages of shared synergies compared to other tests. However, significant differences in temporal patternss were observed between running and all functional tasks, indicating varying activation profiles of shared muscle synergies.

**Conclusion:** Although all functional tests shared muscle synergies with running, variations in the degree of shared synergies and temporal patterns imply that walking, SLH30, and step-up tests may be the most beneficial in predicting running behavior post-ACL injuries. However, functional tests cannot fully replicate running dynamics, suggesting the need for a careful interpretation when assessing readiness for RTR.

## INTRODUCTION

Returning to running (RTR) represents a key milestone in the rehabilitation process following sports-related injuries (1–4). The time frame for RTR can vary greatly; for instance, in the case of anterior cruciate ligament (ACL) injuries, the median time frame for RTR is approximately 12 weeks postoperatively, with a range of 5-39 weeks (1, 2). This phase signifies the transition from early and mid-stage rehabilitation—focused on restoring basic knee function such as regaining adequate knee joint range of motion and resuming everyday activities—to the later stages of rehabilitation. These later stages involve high-intensity training, encompassing activities like jumping, cutting, and sport-specific tasks (5, 6).

Functional tests, which are frequently used as assessment-based criteria, are designed to replicate the physical demands of running. These tests include walk analysis, single leg hops, single leg squats (SLS), and various balance tasks (1, 2). However, it is critical to validate these functional tests as practical benchmarks for facilitating a safe and effective return to running post-injury. This is particularly important for validating tests that share motor control characteristics between running and functional tests.

The role of muscles in motor function is interconnected (7, 8). Essentially, motor control of lower limbs entails the coordinated activation of several lower limb and trunk muscles in response to specific demands requiring both stability and movement. (9). The central nervous system (CNS) is thought to employ an efficient strategy to select the control signal from a large subspace. This is achieved by using a limited set of motor modules or muscle synergies, formed by the flexible combination of muscle activation (10). Thus, exploring the shared muscle synergies between running and commonly used functional tests in clinical settings may provide the neuromechanical rationale for functional tests as RTR criteria following sports-related injuries.

The objective of this study was to determine whether functional tests share muscle synergies with running tasks. Our hypothesis posited that running and functional tests would display shared muscle synergies, but the extent of these synergies would vary between tasks. Additionally, differences in the temporal profiles of these synergies were expected. Such variability could provide critical insights for prioritizing functional tests when evaluating readiness to RTR.

## METHODS

### Participants

Ten healthy males with a mean (SD) age of 21 (±0.3) years old were recruited from the local university. Each participant provided written informed consent for participation in the study. The study was conducted following the principles of the Declaration of Helsinki and approved by the local ethics committee of the University of Tokyo (746).

### Experimental procedures

Participants were asked to freely perform running and nine functional tasks commonly used when considering the return to running following ACL injuries (1), as described in Figure 1. Of note, the average velocities for running and walk were observed to be 2.1 ± 0.28 m/s and 1.2 ± 0.12 m/s, respectively. Each functional test was repeated five times and the order of the tasks was randomly assigned.

**FIGURE 1.**
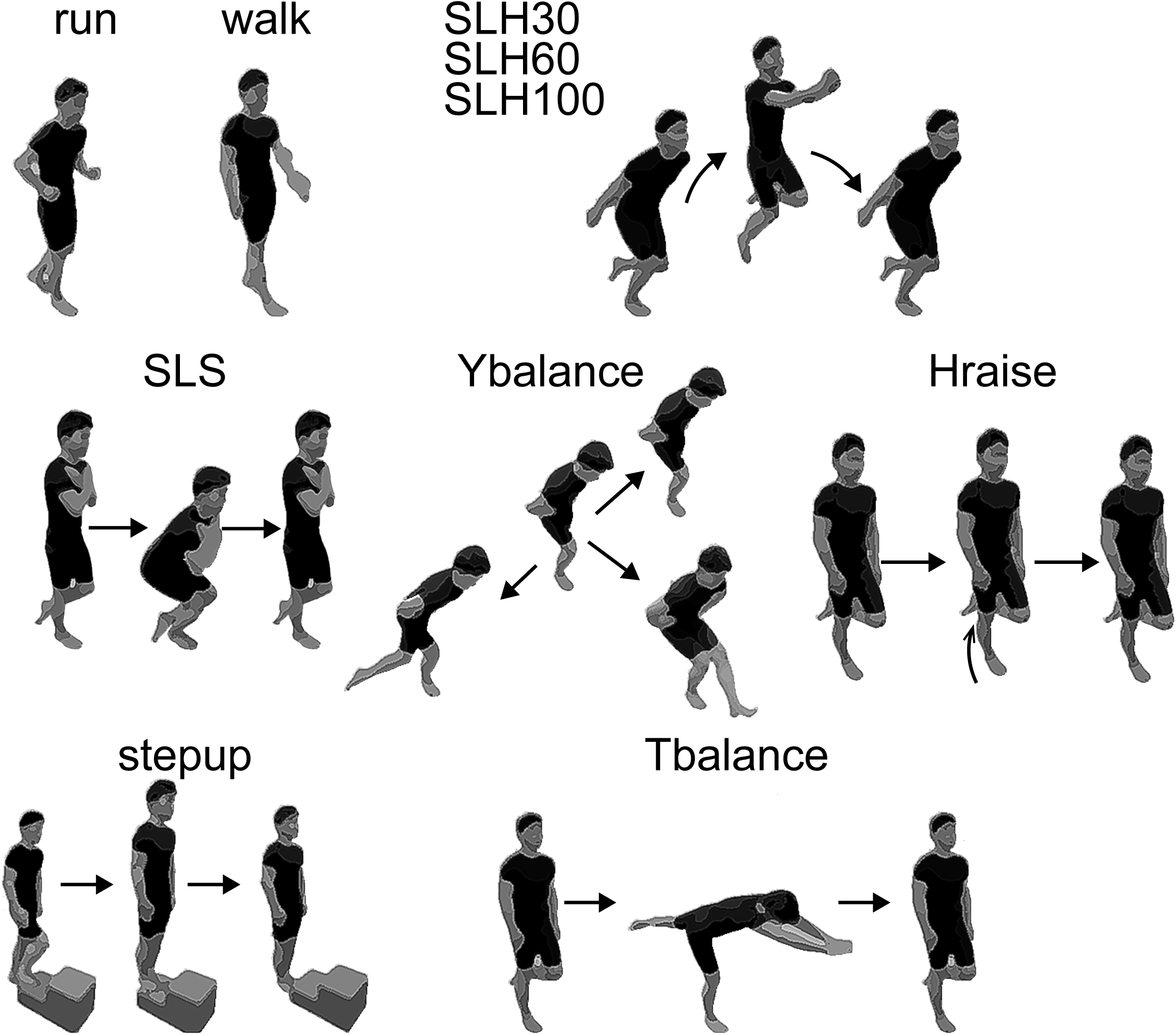
Running and Nine Functional Tests. Participants independently performed running at a jogging pace and a walk. As a result, the average velocities for running and walk were noted to be 2.1 ± 0.28 m/s and 1.2 ± 0.12 m/s, respectively. In the single leg hops (SLH) tests, participants executed a forward jump at 30% (SLH30), 60% (SLH60), and 100% (SLH100) of their maximum distance. A single leg squat (SLS) was carried out with approximately 45 degrees of knee flexion. Hraise refers to the heel raise task, while stepup denotes a forward step up onto a 10 cm height box. Tbalance describes an exercise where the participant stands on the injured leg, forms ac746 ‘T’ shape with the body, drives upward to a standing position, and then slowly returns to the ‘T’ position.

### Data collection

Unilateral surface EMG data were recorded from 16 lower limb and trunk muscle groups: rectus abdominis (RA) (3cm lateral to umbilicus)(11), oblique externus (OE) (15cm lateral to umbilicus)(12), erector spinae at L1 (ESL1) (3cm lateral to the L1 spinous process)(11), gluteus maximus (GM), gluteus medius (Gmed), biceps femoris (long head, BF), semitendinosus (ST), tensor fasciae latae (TFL), adductor longus (ADD), rectus femoris (RF), vastus medialis (VM), vastus lateralis (VL), tibialis anterior (TA), gastrocnemius medialis (MG), soleus (SOL) and peroneus longus (POL). The EMG sensor placements in lower limb were based on SENIAM (surface EMG for a non-invasive assessment of muscles) (13). A wireless EMG system (Trigno Wireless System; DELSYS, Boston, MA, USA) was used to record EMG activity. Each electrode had an inter-electrode spacing of 10 mm. The EMG signals were band-pass filtered (20–450 Hz), amplified (with a 300-gain preamplifier), and sampled at 2000 Hz using an analog-to-digital converter (Power lab/16SP, AD Instruments, Australia).

Marker coordinate data were collected at 120 Hz using an eight-camera motion capture system (Vicon, Centennial, CO) with a 25-marker set. This set incorporated markers for the head, arms, trunk, pelvis, thighs, shanks, and feet, based on the Vicon Plug-in-Gait model. Marker coordinate data were interpolated using cubic spline interpolation to remove gaps in the data and filtered with a low-pass third-order Butterworth filter at 20 Hz. This data was then combined with subject-specific anthropometric data to create an eight-segment whole-body model. The kinematic profiles, calculated by the Vicon Plug-in-Gait model, were used to define the start and end of each trial for each task.

### EMG processing

Raw EMG signals were high-pass filtered at 30 Hz to remove motion artifacts, and then demeaned. The signals were then full-wave-rectified and low-pass filtered at 10 Hz, using a fourth-order Butterworth filter. The smoothed EMG envelopes were time-interpolated to generate 200 time points between the start and end points for each trial so that the EMG data of each trial contributed to the extracted muscle synergies equally.

We created single-task EMG matrices for each task for each participant (that is, the matrix was composed of 16 muscles × 1000 timepoints (the no. of repetitions/cycles (5) × 200 samples)) to extract the muscle synergies for each task. Each EMG from each muscle was normalized to the maximum amplitude across all tasks.

### Independent muscle synergy extraction

To extract muscle synergies, NMF was applied to the single-task EMG matrix. NMF has previously been described as a linear decomposition technique (14, 15) according to equation (1):

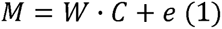

Where *M* (*m*□×□*t* matrix, where *m* is the number of muscles, *t* is the number of samples [i.e., spatiotemporal profiles of muscle activity]) is a linear combination of muscle synergies, *W* (*m*□×lJ*n* matrix, where n is the number of muscle synergies), *C* (*n*□×□*t* matrix, representing temporal patterns), and *e* is the residual error matrix. We applied NMF to extract possible *n* values from 1 to 16 for each dataset. To estimate the optimal number of muscle synergies, the variance accounted for (VAF) by the reconstructed EMG (*M*) was calculated at each iteration (16). VAF was defined as 100 × the square of the uncentered Pearson’s correlation coefficient (16, 17). Considering the local minima inherent in NMF, each synergy extraction was repeated 50 times, and the VAF was calculated at each iteration. Iterations with the highest VAF were maintained (18–21). VAFs > 0.9 were used to identify the optimal number of synergies commonly used in the literature (20–25).

### Shared and specific muscle synergy extraction

To extract the number of shared and running-specific, test-specific muscle synergies, we used a modified version of the NMF algorithm based on the previous studies (24, 26–28) that simultaneously extracts motor modules that are shared across running and each task and those that are specific to each task from a data matrix containing EMG from both conditions. We defined shared synergies as one that activated in both task and thus, temporal pattern components have non-zero coefficients in both tasks. To identify task-specific muscle synergies, the coefficients, C, corresponding to running are set to zero (i.e., test-specific synergies), and to each task are set to zero (i.e., run-specific synergies). Detailed descriptions of this method are in elsewhere (24, 26–28). Briefly, as independent synergy extraction, the number of shared, running specific and task-specific synergies for each participant was determined by the minimum number of total muscle synergies that were required so that the VAF exceeded 90%. We defined the percentage of shared muscle synergies between running and each task as the ratio of the number of shared modules over the number of total motor modules across the two tasks. This modification is thought to improve the accuracy shared and task-specific muscle synergies by minimizing the possibility in which the numbers of synergies are underestimated when synergies are extracted from the EMG of each task independently, and compare the similarity of synergies between tasks to identify shared and task-specific synergies (26, 29).

We identified representative shared, running-specific and test-specific synergies across participants using hierarchical clustering analysis (Ward’s method, Euclidian distance) of muscle weighting components (19). To determine the optimal number of clusters, we computed the gap statistic (30), which measures the compactness of the clustering achieved against those in reference data sets without any obvious clustering similar to a previous study (31). Reference data sets (N□=□500) were initially generated by sampling uniformly from within the bounds of the original muscle-synergy set; each of them was then clustered by the hierarchical cluster, at 2–20 clusters. The optimal number of clusters was then the smallest number, h, such that

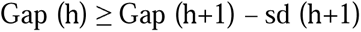

where Gap(k) represents the gap statistic at h clusters, and sd(h) signifies the standard deviation of the clustering compactness within the reference data sets (30).

We defined shared, running-specific and test-specific synergy clusters as having each three types of synergies from ≥ 1/2 of synergies within a cluster. If the cluster was not contributed by any types of synergies, we defined it as “none”.

### Statistics

We compared the percentage of shared muscle synergies between tasks. The values were compared using the Friedman test, which is a non-parametric method for multiple comparisons of independent samples, as a normal distribution was not observed in the data (tested using the Shapiro–Wilk test). When the Friedman test showed significant effects, multiple comparison post-hoc analyses were performed using the Wilcoxon signed-rank test.

Temporal pattern components were compared between running and functional tests using statistical parametric mapping (SPM) (spm1d v0.4.7 for MATLAB, Institute of Neurology, London, UK) (32). Since each single-task EMG matrix contained 5 repetitions/cycles, the extracted temporal pattern components of running and functional tests were converted into an averaged repetition/cycles for each participant before comparing temporal pattern components using SPM analysis.

The p values obtained from were corrected using the false discovery rate (FDR) correction for multiple comparisons (33). The significance level for all tests was set at *p*□<□0.05. When there was a significant difference between the groups, effect sizes (ES) were calculated using Cohen’s d (34). We recruited ten participants without an a priori power analysis, thus, we instead conducted a sensitivity analysis in G*Power, which indicated that an effect size of 0.71 would be necessary to obtain a power of 80% at an α of 0.05.

## RESULTS

### Muscles synergies for running and functional tests

Table 1 presents the VAF values between the groups for running and functional tests. Figure 2 depicts the percentage of muscle synergies shared between running and each respective functional test. The median percentage of shared synergies for each test were as follows: walk (75% [40%-100%]), SLH30 (60% [20%-100%]), SLH60 (50% [28%-75%]), SLH100 (50% [28%-80%]), SLS (32% [16%-50%]), Ybalance (40% [25%-80%]), Hraise (22% [20%-60%]), stepup (63% [0%-100%]), and Tbalance (40% [20%-80%]). The percentage of shared synergies was significantly different among the nine representative speeds in non-runners and runners (*p* = 0.000079 for both groups, Friedman test one-way ANOVA). Statistically significant differences were observed between the following pairings: Walk and SLS (p = 0.0038, ES = 2.18), Walk and Hraise (p = 0.0098, ES = 2.05), SLH30 and SLS (p =0.0017, ES = 1.56), SLH30 and Hraise (p = 0.0039, ES = 1.47), SLH and stepup (p = 0.0072, ES = 1.31), and Hraise and stepup (p = 0.0176, ES = 1.25).

**FIGURE 2.**
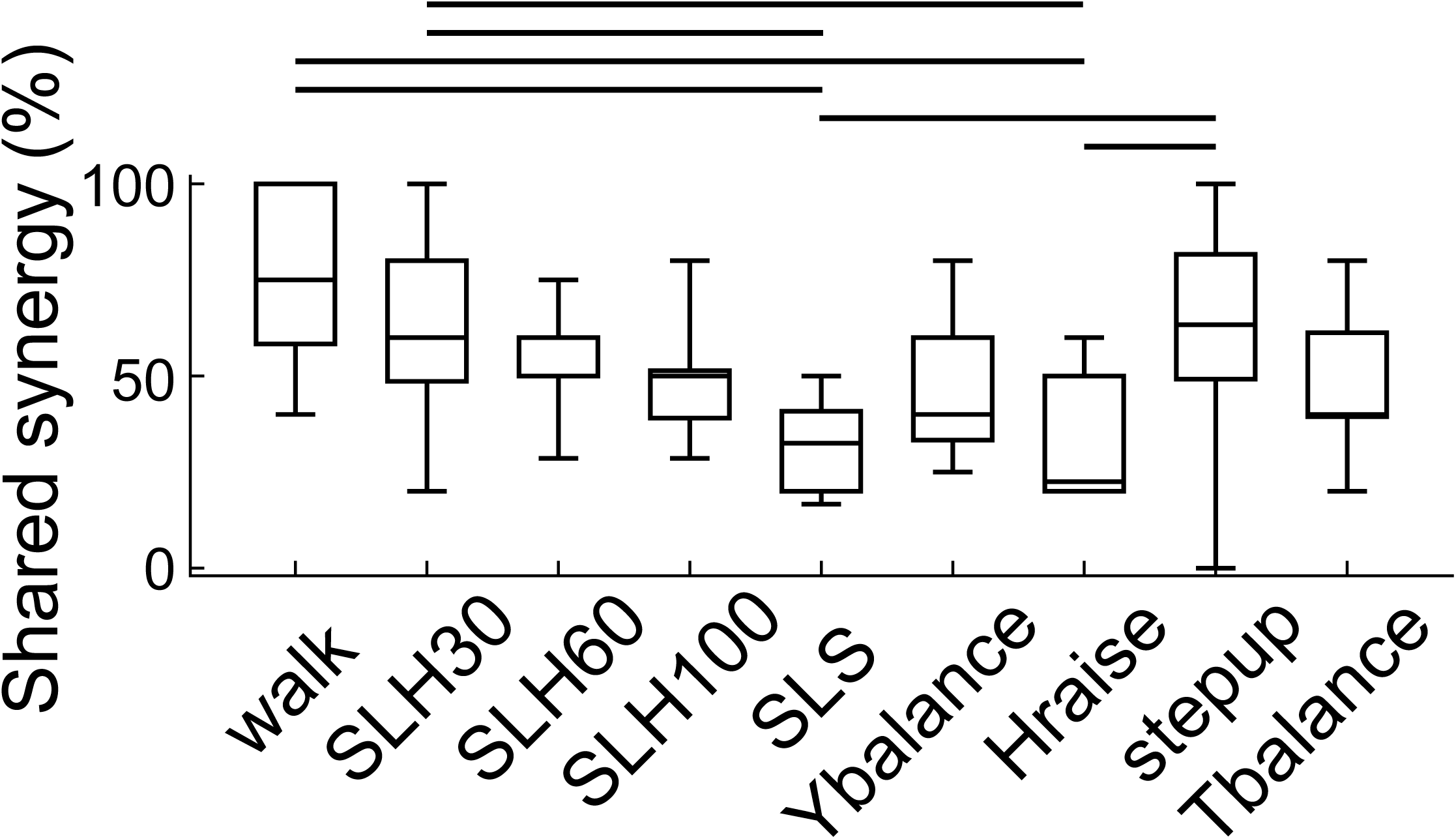
The percent of shared muscle synergies between running and each functional test. Median values are indicated as horizontal lines inside the boxes. The edges of the boxes represent the 25th and 75th percentiles. (*p* = 0.000079 for both groups, Friedman test one-way ANOVA). Statistically significant differences were observed between the following pairings: Walk and SLS (p = 0.0038, ES = 2.18), Walk and Hraise (p = 0.0098, ES = 2.05), SLH30 and SLS (p =0.0017, ES = 1.56), SLH30 and Hraise (p = 0.0039, ES = 1.47), SLH and stepup (p = 0.0072, ES = 1.31), and Hraise and stepup (p = 0.0176, ES =1.25).

**TABLE 1.** Variance account for (VAF) and number of muscle synergies for running and functional tests

|  |  | VAF values | Number of synergies |
| --- | --- | --- | --- |
| Running | | 0.92 (± 0.01) | 3.9 (± 0.31) |
| Functional tests | Walk | 0.92 (± 0.01) | 3.8 (± 0.63) |
| | SLH30 | 0.92 (± 0.01) | 4.3 (± 0.67) |
| | SLH60 | 0.91 (± 0.01) | 4.6 (± 0.51) |
| | SLH90 | 0.91 (± 0.01) | 5.1 (± 0.56) |
| | SLS | 0.92 (± 0.01) | 2.4 (± 0.51) |
| | Ybalance | 0.92 (± 0.01) | 3.6 (± 0.84) |
| | Hraiese | 0.92 (± 0.01) | 2.3 (± 0.67) |
| | Steuup | 0.91 (± 0.01) | 4.5 (± 0.97) |
| | Tbalance | 0.92 (± 0.01) | 3.4 (± 0.69) |

Figure 3 shows representative shared, running-specific, and test-specific muscle synergies identified by cluster analysis (muscle weighting components (W) and temporal pattern components (C)). All functional tests shared the muscle synergies with running although the number of shared synergies differed between functional tests. We also identified the running-specific and test-specific muscle synergies in all functional tests except walk task. The SPM analysis found significant differences in temporal pattern components between running and each functional task (p < 0.05).

**FIGURE 3.**
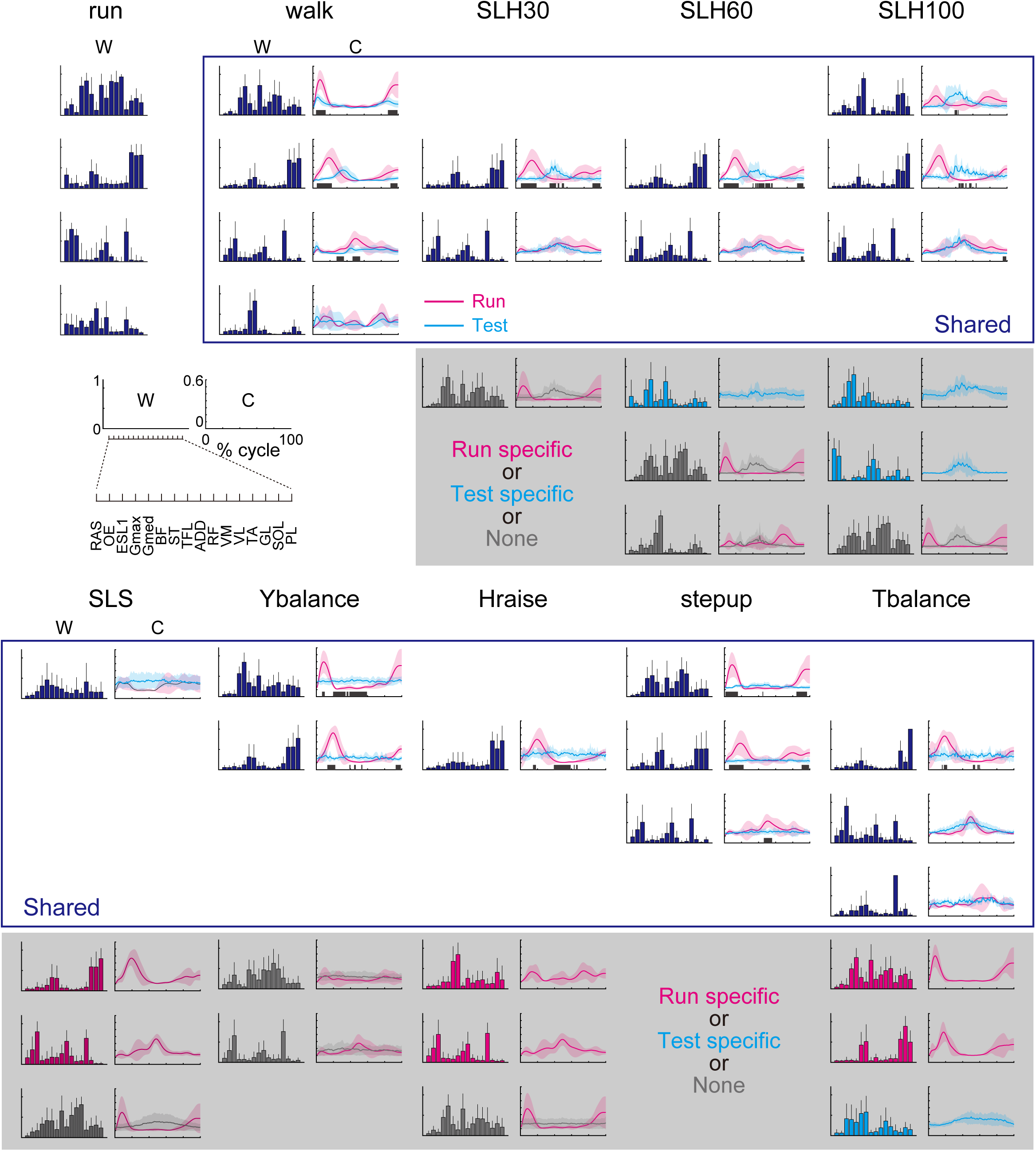
Representative shared, running-specific, and test-specific muscle synergies identified by cluster analysis (muscle weighting components (W) and temporal pattern components (C)). The gray sections on the X-axis of C represent the temporal pattern components where a significant difference between running and functional tasks was observed (p < 0.05). RA: rectus abdominis, OE: oblique externus, ESL1(erector spinae at L1), GM: gluteus maximus, Gmed: gluteus medius, BF: biceps femoris, ST: semitendinosus, TFL: tensor fasciae latae, ADD: adductor longus, RF: rectus femoris, VM: vastus medialis, VL: vastus lateralis, TA: tibialis anterior, MG: gastrocnemius medialis, SOL: Soleus (SOL) and POL: peroneus longus.

## DISCUSSION

In this study, we applied the NMF algorithm to large-scale and high-dimensional EMG data to investigate shared muscle synergies between running and functional tasks that are commonly used to determine when individuals with ACL injuries can return to running activities. Overall, although all functional tests shared muscle synergies with running, our results suggest that walking, SLH, and step-up tests could be the most beneficial because the percentages of shared synergies between running and these tests were significantly higher than those with other functional tests. However, despite these shared muscle synergies, there were notable differences in the temporal patterns between running and functional tasks. These discrepancies suggest the need for caution when using functional tests alone to predict running capabilities post-injury.

We hypothesized that a high degree of shared synergies between running and functional tests would signify these tests as strong predictors of running behavior, due to similar motor control characteristics, including the coordination of multiple muscles. Firstly, our results showed that the percentage of shared synergies (muscle weighting components) between running and walking was 75% [40%-100%], with all synergies between the two tasks being shared. This aligns with a previous study that suggested consistent muscle weighting components for walking and running, facilitating the transition from walking to running. (35, 36). Similarly, we found that the step-up test shared a significant amount of muscle synergies with running, which is reasonable given that both activities share the same mechanical goal of propelling the body’s center of mass forward. (37–39). This action requires dynamic balance with appropriate activations of lower and trunk muscles. The SLH test, particularly the SLH30 variant, exhibited a higher degree of shared muscle synergies with running compared to other non-jumping tests. Conversely, SLH60 and SLH100 demonstrated a relatively low number of shared synergies, as they also generated test-specific synergies beneficial for longer forward jumping.

NMF also extracted temporal pattern components, signifying the activation profiles of muscle weighting components. Notably, there were significant differences in some temporal pattern components of muscle synergies between running and all functional tests. From a neuromechanical perspective, these temporal patterns reflect the specific timing and intensity of muscle activation, illustrating that even though muscle synergies may be similar, the way they are employed in different activities can differ significantly. This discrepancy could be attributed to the unique demands of each task. Even though similar muscle groups may be engaged (thus the shared synergies), the coordination, timing, and intensity of muscle activation might not perfectly align. (40, 41). From a clinical perspective, these differences suggest that functional tests cannot fully replicate running behavior. Specifically, the characteristics of temporal components of walking, SLH30, and stepup tasks that exhibited higher percentages of shared synergies, differed significantly compared to those in running. These disparities in temporal patterns, despite shared muscle synergies, can limit the interpretation of the results, suggesting that the differing activation profiles of shared muscle synergies in these functional tests may impede their predictability of safe running performance. Clinicians should consider that while these functional tests can help assess readiness for running by reflecting certain shared muscle activation patterns, they do not perfectly mimic the exact dynamics of running. Therefore, clinicians should utilize functional tests in combination with running evaluations in a clinical setting using a treadmill whenever possible. Additionally, other assessments such as knee range of motion, strength, and psychological readiness for running should be taken into account. A comprehensive assessment will aid in detecting basic function for running and deviations from normal running patterns before athletes resume running outside a clinical setting (1, 2).

Steady progress through high-quality rehabilitation is essential for functional recovery (42). Resuming sports activities prematurely can elevate the risk of secondary injuries (43), while excessively slow progress might adversely affect motivation and psychological preparedness for sports performances (44). Consequently, determining the appropriate timing for a return to running (RTR) is a crucial milestone in the effective rehabilitation continuum for a return to sports (1, 2). Our findings provide a novel rationale for using functional tests in decision-making for RTR. Further exploration into the degree of shared synergies between running and functional tests among individuals with and without lower-limb sports injuries could yield intriguing results. This, along with investigating the relationship between the degree of shared synergies and future injuries, could potentially offer valuable biomarkers for injury prevention.

Note of caution in interpreting the study findings is warranted. Although numerous functional tests mimic the motor control patterns or muscle synergies of running, the running speed employed in this study was relatively low, averaging 2.1 ± 0.28 m/s. This raises uncertainty regarding whether the functional tests used also capture the muscle synergy variations of high-speed running, direction changes, and cutting movements, all of which are high-risk activities for lower limb sports injuries. Indeed, a previous study highlighted distinct muscle synergies at different speeds (19), If the tests used don’t replicate these running variations, we might not accurately predict running ability and may need to consider other functional test batteries.

## CONCLUSION

Our study suggests that the walk, SLH, and step-up tests can be reliable indicators of running behavior due to their shared muscle synergies with running. Despite different temporal pattern components in these tests compared to running, they offer a practical means to assess running ability. However, clinicians should be aware that these functional tests may not fully emulate the physical demands of running. Therefore, these tests should be incorporated as key components in the comprehensive decision-making process for a return to running.

## Declarations

### Ethics approval and consent to participate

All experimental protocols were approved by the local ethics committee of the University of Tokyo, and all participants gave their written informed consent.

### Consent for publication

The participant depicted in the photos gave their written consent for publication.

### Availability of data and materials

The datasets used and/or analyzed during the current study are available from the corresponding author on reasonable request.

## Acknowledgements

Not applicable.

## References

1. Rambaud AJM, Ardern CL, Thoreux P, Regnaux JP, Edouard P. Criteria for return to running after anterior cruciate ligament reconstruction: a scoping review. British journal of sports medicine. 2018;52(22):1437–44. Epub 20180502. doi: 10.1136/bjsports-2017-098602. PubMed PMID: 29720478.

2. Van Cant J, Pairot de Fontenay B, Douaihy C, Rambaud A. Characteristics of return to running programs following an anterior cruciate ligament reconstruction: A scoping review of 64 studies with clinical perspectives. Phys Ther Sport. 2022;57:61–70. Epub 20220719. doi: 10.1016/j.ptsp.2022.07.006. PubMed PMID: 35921783.

3. Macdonald B, McAleer S, Kelly S, Chakraverty R, Johnston M, Pollock N. Hamstring rehabilitation in elite track and field athletes: applying the British Athletics Muscle Injury Classification in clinical practice. British journal of sports medicine. 2019;53(23):1464–73. Epub 20190712. doi: 10.1136/bjsports-2017-098971. PubMed PMID: 31300391.

4. Vergani L, Cuniberti M, Zanovello M, Maffei D, Farooq A, Eirale C. Return to Play in Long-Standing Adductor-Related Groin Pain: A Delphi Study Among Experts. Sports Med Open. 2022;8(1):11. Epub 20220118. doi: 10.1186/s40798-021-00400-z. PubMed PMID: 35043267; PubMed Central PMCID: PMC8766680.

5. Myer GD, Ford KR, McLean SG, Hewett TE. The effects of plyometric versus dynamic stabilization and balance training on lower extremity biomechanics. Am J Sports Med. 2006;34(3):445–55.

6. Buckthorpe M, Della Villa F. Optimising the ‘Mid-Stage’ Training and Testing Process After ACL Reconstruction. Sports medicine (Auckland, NZ). 2020;50(4):657–78. doi: 10.1007/s40279-019-01222-6. PubMed PMID: 31782065.

7. Dickinson MH, Farley CT, Full RJ, Koehl MA, Kram R, Lehman S. How animals move: an integrative view. Science. 2000;288(5463):100–6. doi: 10.1126/science.288.5463.100. PubMed PMID: 10753108.

8. Chiel HJ, Ting LH, Ekeberg O, Hartmann MJ. The brain in its body: motor control and sensing in a biomechanical context. J Neurosci. 2009;29(41):12807–14. doi: 10.1523/JNEUROSCI.3338-09.2009. PubMed PMID: 19828793; PubMed Central PMCID: PMC2794418.

9. Ting LH, Chiel HJ, Trumbower RD, Allen JL, McKay JL, Hackney ME, et al. Neuromechanical principles underlying movement modularity and their implications for rehabilitation. Neuron. 2015;86(1):38–54. doi: 10.1016/j.neuron.2015.02.042.

10. d’Avella A, Portone A, Fernandez L, Lacquaniti F. Control of fast-reaching movements by muscle synergy combinations. J Neurosci. 2006;26(30):7791–810. doi: 10.1523/JNEUROSCI.0830-06.2006. PubMed PMID: 16870725; PubMed Central PMCID: PMC6674215.

11. McGill S, Juker D, Kropf P. Appropriately placed surface EMG electrodes reflect deep muscle activity (psoas, quadratus lumborum, abdominal wall) in the lumbar spine. Journal of biomechanics. 1996;29(11):1503–7. doi: 10.1016/0021-9290(96)84547-7. PubMed PMID: 8894932.

12. Vera-Garcia FJ, Moreside JM, McGill SM. Abdominal muscle activation changes if the purpose is to control pelvis motion or thorax motion. Journal of electromyography and kinesiology : official journal of the International Society of Electrophysiological Kinesiology. 2011;21(6):893–903. Epub 20110916. doi: 10.1016/j.jelekin.2011.08.003. PubMed PMID: 21925900.

13. Hermens HJ, Freriks B, Disselhorst-Klug C, Rau G. Development of recommendations for SEMG sensors and sensor placement procedures. Journal of electromyography and kinesiology : official journal of the International Society of Electrophysiological Kinesiology. 2000;10(5):361–74. doi: 10.1016/s1050-6411(00)00027-4. PubMed PMID: 11018445.

14. Lee DD, Seung HS. Learning the parts of objects by non-negative matrix factorization. Nature. 1999;401(6755):788–91. doi: 10.1038/44565.

15. Tresch MC, Cheung VCK, d’Avella A. Matrix factorization algorithms for the identification of muscle synergies: evaluation on simulated and experimental data sets. J Neurophysiol. 2006;95(4):2199-212. doi: 10.1152/jn.00222.2005.

16. Torres-Oviedo G, Macpherson JM, Ting LH. Muscle synergy organization is robust across a variety of postural perturbations. J Neurophysiol. 2006;96(3):1530–46. doi: 10.1152/jn.00810.2005.

17. Zar JH. Biostatistical analysis: Pearson Education India; 1999 1999.

18. Yokoyama H, Kato T, Kaneko N, Kobayashi H, Hoshino M, Kokubun T, et al. Basic locomotor muscle synergies used in land walking are finely tuned during underwater walking. Sci Rep. 2021;11(1). doi: 10.1038/s41598-021-98022-8.

19. Yokoyama H, Ogawa T, Kawashima N, Shinya M, Nakazawa K. Distinct sets of locomotor modules control the speed and modes of human locomotion. Sci Rep. 2016;6:36275. doi: 10.1038/srep36275. PubMed Central PMCID: PMC5090253.

20. Saito H, Yokoyama H, Sasaki A, Kato T, Nakazawa K. Evidence for basic units of upper limb muscle synergies underlying a variety of complex human manipulations. J Neurophysiol. 2022;127(4):958–68. Epub 20220302. doi: 10.1152/jn.00499.2021. PubMed PMID: 35235466.

21. Saito H, Yokoyama H, Sasaki A, Kato T, Nakazawa K. Flexible Recruitments of Fundamental Muscle Synergies in the Trunk and Lower Limbs for Highly Variable Movements and Postures. Sensors. 2021;21(18). doi: 10.3390/s21186186.

22. Botzheim L, Laczko J, Torricelli D, Mravcsik M, Pons JL, Oliveira Barroso F. Effects of gravity and kinematic constraints on muscle synergies in arm cycling. J Neurophysiol. 2021;125(4):1367–81. doi: 10.1152/jn.00415.2020.

23. Stamenkovic A, Ting LH, Stapley PJ. Evidence for constancy in the modularity of trunk muscle activity preceding reaching: Implications for the role of preparatory postural activity. J Neurophysiol. 2021. doi: 10.1152/jn.00093.2021.

24. Sawers A, Allen JL, Ting LH. Long-term training modifies the modular structure and organization of walking balance control. J Neurophysiol. 2015;114(6):3359–73. doi: 10.1152/jn.00758.2015. PubMed Central PMCID: PMC4868379.

25. Hug F, Turpin NA, Couturier A, Dorel S. Consistency of muscle synergies during pedaling across different mechanical constraints. J Neurophysiol. 2011;106(1):91–103. doi: 10.1152/jn.01096.2010.

26. Cheung VCK, d’Avella A, Tresch MC, Bizzi E. Central and sensory contributions to the activation and organization of muscle synergies during natural motor behaviors. J Neurosci. 2005;25(27):6419–34. doi: 10.1523/JNEUROSCI.4904-04.2005. PubMed Central PMCID: PMC6725265.

27. Cheung VCK, d’Avella A, Bizzi E. Adjustments of motor pattern for load compensation via modulated activations of muscle synergies during natural behaviors. J Neurophysiol. 2009;101(3):1235–57. doi: 10.1152/jn.01387.2007. PubMed Central PMCID: PMC2666413.

28. Allen JL, McKay JL, Sawers A, Hackney ME, Ting LH. Increased neuromuscular consistency in gait and balance after partnered, dance-based rehabilitation in Parkinson’s disease. J Neurophysiol. 2017;118(1):363–73. Epub 20170405. doi: 10.1152/jn.00813.2016. PubMed PMID: 28381488; PubMed Central PMCID: PMC5501921.

29. Cheung VC, d’Avella A, Bizzi E. Adjustments of motor pattern for load compensation via modulated activations of muscle synergies during natural behaviors. J Neurophysiol. 2009;101(3):1235–57. Epub 20081217. doi: 10.1152/jn.01387.2007. PubMed PMID: 19091930; PubMed Central PMCID: PMC2666413.

30. Tibshirani R, Walther G, Hastie T. Estimating the number of clusters in a data set via the gap statistic. J R Stat Soc Series B Stat Methodol. 2001;63(2):411-23. doi: 10.1111/1467-9868.00293.

31. Cheung VCK, Cheung BMF, Zhang JH, Chan ZYS, Ha SCW, Chen C-Y, et al. Plasticity of muscle synergies through fractionation and merging during development and training of human runners. Nat Commun. 2020;11(1):4356. doi: 10.1038/s41467-020-18210-4. PubMed Central PMCID: PMC7459346.

32. Pataky TC. Generalized n-dimensional biomechanical field analysis using statistical parametric mapping. Journal of biomechanics. 2010;43(10):1976–82. doi: 10.1016/j.jbiomech.2010.03.008. PubMed PMID: 20434726.

33. Benjamini Y, Hochberg Y. Controlling the False Discovery Rate: A Practical and Powerful Approach to Multiple Testing. Journal of the Royal Statistical Society: Series B (Methodological). 1995;57(1):289–300. doi: https://doi.org/10.1111/j.2517-6161.1995.tb02031.x.

34. Cohen J. Statistical power analysis for the behavioral sciences. Abingdon. England: Routledge. 1988.

35. Cappellini G, Ivanenko YP, Poppele RE, Lacquaniti F. Motor Patterns in Human Walking and Running. Journal of Neurophysiology. 2006;95(6):3426–37. doi: 10.1152/jn.00081.2006.

36. Hagio S, Fukuda M, Kouzaki M. Identification of muscle synergies associated with gait transition in humans. Front Hum Neurosci. 2015;9:48. doi: 10.3389/fnhum.2015.00048. PubMed Central PMCID: PMC4322718.

37. Cook TM, Zimmermann CL, Lux KM, Neubrand CM, Nicholson TD. EMG Comparison of Lateral Step-up and Stepping Machine Exercise. J Orthop Sports Phys Ther. 1992;16(3):108–13. doi: 10.2519/jospt.1992.16.3.108. PubMed PMID: 18796761.

38. Simenz CJ, Garceau LR, Lutsch BN, Suchomel TJ, Ebben WP. Electromyographical analysis of lower extremity muscle activation during variations of the loaded step-up exercise. Journal of strength and conditioning research / National Strength & Conditioning Association. 2012;26(12):3398–405. doi: 10.1519/JSC.0b013e3182472fad. PubMed PMID: 22237139.

39. Wang MY, Flanagan S, Song JE, Greendale GA, Salem GJ. Lower-extremity biomechanics during forward and lateral stepping activities in older adults. Clin Biomech (Bristol, Avon). 2003;18(3):214–21. doi: 10.1016/s0268-0033(02)00204-8. PubMed PMID: 12620784; PubMed Central PMCID: PMC3460801.

40. d’Avella A, Bizzi E. Shared and specific muscle synergies in natural motor behaviors. Proc Natl Acad Sci U S A. 2005;102(8):3076–81. doi: 10.1073/pnas.0500199102. PubMed Central PMCID: PMC549495.

41. Bizzi E, Cheung VCK. The neural origin of muscle synergies. Front Comput Neurosci. 2013;7:51. doi: 10.3389/fncom.2013.00051. PubMed Central PMCID: PMC3638124.

42. Grindem H, Granan LP, Risberg MA, Engebretsen L, Snyder-Mackler L, Eitzen I. How does a combined preoperative and postoperative rehabilitation programme influence the outcome of ACL reconstruction 2 years after surgery? A comparison between patients in the Delaware-Oslo ACL Cohort and the Norwegian National Knee Ligament Registry. British journal of sports medicine. 2015;49(6):385–9. Epub 20141028. doi: 10.1136/bjsports-2014-093891. PubMed PMID: 25351782; PubMed Central PMCID: PMC4351141.

43. Kyritsis P, Bahr R, Landreau P, Miladi R, Witvrouw E. Likelihood of ACL graft rupture: not meeting six clinical discharge criteria before return to sport is associated with a four times greater risk of rupture. Br J Sports Med. 2016;50(15):946–51. doi: 10.1136/bjsports-2015-095908.

44. Ardern CL, Taylor NF, Feller JA, Whitehead TS, Webster KE. Psychological responses matter in returning to preinjury level of sport after anterior cruciate ligament reconstruction surgery. Am J Sports Med. 2013;41(7):1549–58. Epub 20130603. doi: 10.1177/0363546513489284. PubMed PMID: 23733635.

